# Recent hybridisation and ghost introgression among a trio of island passerines

**DOI:** 10.64898/2026.05.06.723294

**Authors:** Abby Williams, Andrea Estandía, Ashley T. Sendell-Price, Amanda M. Carpenter, Dmitry Filatov, Kristen Ruegg, Sonya M. Clegg

## Abstract

Hybridisation between species was once considered a relatively uncommon occurrence but is now recognised to occur frequently across many different taxa. It can result in homogenisation of previously distinct forms, a potential conservation issue, but can also act as a catalyst for diversification through introgression and sharing of favourable genes. Repeated rounds of island colonisation followed by speciation result in secondary sympatry, with the potential for hybridisation between early and late arrivers. In the southwest Pacific, this situation has arisen in the avian family Zosteropidae (the white-eyes). Here we use whole genome sequencing of live birds and historical specimens to characterise hybridisation between three white-eye species on Norfolk Island: two island endemics, *Zosterops tenuirostris* and the now-extinct *Zosterops albogularis*, and *Zosterops lateralis*, which colonised the island in 1904. Despite over two million years of divergence between *Z. lateralis* and the two endemics, we provide genomic evidence of their hybridisation. First, we confirm the identities of three *Z. lateralis x Z. tenuirostris* hybrids and additionally identify one *Z. lateralis x Z. albogularis* hybrid. We also report asymmetric, genome-wide introgression from both endemics into *Z. lateralis*, with introgressed regions enriched for a range of potential functions. However, despite this introgression, species boundaries have been maintained, and the extant endemic *Z. tenuirostris* does not appear to be at risk of genetic extinction. Our work additionally demonstrates an unusual case of recent ghost introgression from the extinct *Z. albogularis* into *Z. lateralis*. This study sheds light on the genomic outcomes of secondary sympatry and its potential consequences for single-island endemics.

## Introduction

Hybridisation, a phenomenon once considered to be atypical in nature (Mayr, 1943), is now known to be taxonomically and geographically widespread (Ellegren et al., 2012; Meier et al., 2017; Ottenburghs, 2023; Payseur & Rieseberg, 2016; Sankararaman et al., 2014; Taylor & Larson, 2019; Tiesmeyer et al., 2020). The evolutionary outcomes of hybridisation and its impact on biodiversity patterns are varied (Cowles & Uy, 2019; Moran et al., 2021). At one extreme, pervasive hybridisation can homogenise biodiversity by collapsing formerly distinct lineages or species into a single hybrid lineage with an admixed genome (Todesco et al., 2016; Webb et al., 2011). At the other extreme, exclusive breeding among hybrid individuals can generate a new population of mixed ancestry that remains distinct from both parental populations, facilitating hybrid speciation (Hermansen et al., 2011; Lamichhaney et al., 2018; Larsen et al., 2010). Beyond these exceptional ways to homogenise or generate biodiversity, introgression - hybridisation combined with backcrossing of hybrid individuals into parental populations - provides a mechanism for transfer of genetic variation from one species to another, without compromising the integrity of parental species (Harrison & Larson, 2014). As has been demonstrated in an increasing number of studies, introgression can be an important source of novel variation on which natural selection can act (Gaczorek et al., 2024; Meier et al., 2017; Valencia-Montoya et al., 2020; Wu et al., 2018).

Secondary contact between species that initially diverged in allopatry provides an opportunity for hybridisation, with the outcome depending on the characteristics of the species involved. Species that have diverged sufficiently in allopatry may hybridise only as a last resort due to restricted mate choice in secondary contact, resulting in a pulse of hybridisation that ceases soon after contact (Hubbs, 1955; Moran et al., 2021; Sardell & Uy, 2016). Alternatively, less divergent species may hybridise extensively, not yet having evolved barriers to reproduction (Cowles & Uy, 2019). In this way, secondary contact has been suggested to act as a sorting process, only permitting those species that have diverged sufficiently in allopatry to coexist in sympatry (Anderson & Matute, 2025).

The dynamics of secondary sympatry depend on the demographic, geographic, and ecological conditions under which secondary contact is established. Studies of secondary contact have typically focused on systems where sympatry is established due to an expanding population front on a large land mass, which is likely to have different population dynamics to that arising from a rare colonisation event of a geographically isolated island. In the former, it may frequently be the case that both colonising and incumbent forms are relatively abundant (e.g. Garroway et al., 2010; Jansson et al., 2007), or the incumbent form is rarer (e.g. Smith et al., 2022; Tiesmeyer et al., 2020) and that the colonising species is demographically and genetically well connected to other parts of its distribution. In contrast, where a small number of individuals colonise an isolated island, the colonising species is likely to be rarer, at least initially, and separated from its source population where the colonising event results from infrequent long-distance dispersal (Reatini & Vision, 2024).

This imbalance in numbers between native and colonising species could promote hybridisation through restricting mate choice of the coloniser - the “desperation hypothesis” (Hubbs, 1955; McCracken & Wilson, 2011). Cases of recently formed secondary sympatry resulting from naturally occurring island colonisation events are rare, and few have been studied from a genetic perspective. Shogren et al. (2024) reported ongoing, asymmetric mtDNA and nuclear DNA introgression between two species of *Myzomela* honeyeaters in the Solomon Islands that formed sympatric populations in the late 19th to early 20th century, and captured a number of phenotypic hybrids. Similarly, in the Galapagos islands, the documented arrival of *Geospiza conirostris* in 1981 from Española on Daphne Major led to hybridisation with a native congener, ultimately resulting in a hybrid speciation event (Lamichhaney et al., 2018). Despite these isolated examples, the processes and evolutionary consequences of secondary sympatry on islands remain poorly understood, and further studies are needed to establish general principles governing its outcomes.

The white-eyes (family Zosteropidae) are an example of a geographic radiation that has rapidly evolved the vast majority of its 100+ species within the last two million years (Moyle et al., 2009). Members of this family are widely distributed across Africa, the Asian subcontinent, and Australia, with many parapatric and sympatric occurrences (Mees, 1969; Winkler et al., 2020). White-eyes have also been highly successful at colonising islands, especially those of the Indian and Pacific Oceans (Mees, 1969). There are many examples where independent colonisations have resulted in sympatric distributions on islands e.g., Sri Lanka (two species) (Wickramasinghe et al., 2017) Kolombangara, Solomon Islands (two species) (Cowles & Uy, 2019), and islands in the Indonesian archipelago (up to four species) (Gwee et al., 2020). Despite numerous opportunities provided by secondary sympatry on both continents and islands, and their relatively recent diversification (Moyle et al., 2009), members of the family do not appear to frequently hybridise (Ottenburghs et al., 2015).

Hybridisation has been suggested to occur between sympatric species in southern Africa (Oatley et al., 2017) and continental Australia (Degnan, 1993), but not in East Africa cases (Husemann et al., 2014) or the Solomon Islands in the Pacific (Cowles & Uy, 2019). It has been suggested that the rapid evolution of reproductive isolation in Zosteropidae supports the maintenance of species boundaries upon secondary contact, contributing to species diversity in the family (Cowles & Uy, 2019). Taken together, this evidence raises questions about the role of hybridisation in the diversification of the Zosteropidae.

In the southwest Pacific, the silvereye (*Zosterops lateralis*) has repeatedly colonised isolated islands and archipelagos over the last million years, either directly from the Australian continent or via island-hopping (Estandía et al., 2025). This has resulted in numerous sympatric *Zosterops* species pairs or triplets; for example, on Lifou, New Caledonia, *Z. lateralis* co-occurs with two single-island endemics (Mees, 1969), and across the Vanuatu archipelago, *Z. lateralis* co-occurs with the endemic *Z. flavifrons* on 11 of 13 main islands (Clegg & Phillimore, 2010). A more recent case of sympatry occurred following the sequential colonisation of the Tasmanian silvereye subspecies (*Z. l. lateralis, Z. lateralis* hereafter) to Norfolk Island, via New Zealand (North, 1904; Fig. 1A). When *Z. lateralis* arrived on this geographically remote oceanic island in 1904, it formed a sympatric triplet with two endemic *Zosterops* species - the slender-billed white-eye (*Z. tenuirostris*) and the white-chested white-eye (*Z. albogularis;* Fig. 1B). The two endemics diverged in allopatry from *Z. lateralis,* sharing a most recent common ancestor ∼ 2 mya (Estandía et al., 2025).

**Figure 1.**
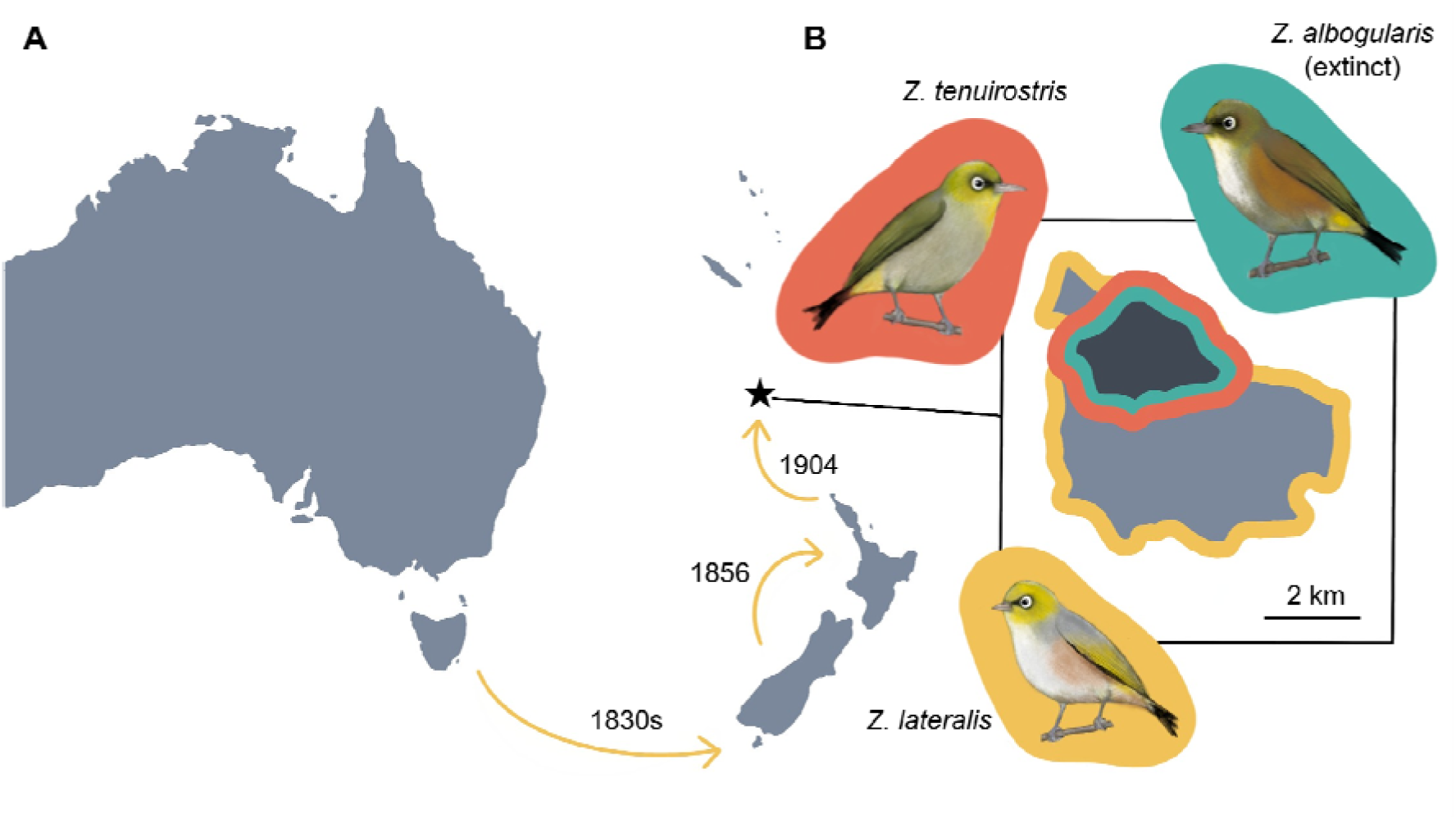
A) Colonisation history of *Z. lateralis*, from Tasmania (Australia) to the South Island (New Zealand) in the 1830s, then to the North Island (New Zealand) in 1856, and finally to Norfolk Island (black star) in 1904. B) Map of Norfolk Island showing boundaries of Norfolk Island National Park (black), where *Z. tenuirostris* is mainly found today, though both endemics likely occurred throughout the island when it was still forested. *Z. lateralis* has an island-wide distribution. Bird illustrations by AW.

Upon colonisation, the *Z. lateralis* population rapidly increased, becoming more common than either of the endemics by 1910 (Mees, 1969). It has persisted as a large population (likely many thousands of individuals) and is now the most abundant *Zosterops* species in both native forests and disturbed habitats (Dutson, 2013). *Z. tenuirostris* is classified as near threatened due to its restricted insular distribution (Garnett et al., 2011) though as of 2013 its population size was estimated as at least several thousand individuals (Dutson, 2013).

However, it was no longer the most abundant forest bird (as it was in a 1962 survey; Mees, 1969), with some evidence of decline since 2000, though ongoing surveys are needed to assess long-term trends (Dutson, 2013). The oldest endemic, *Z. albogularis*, was still common in the first decade of the 20th century (Hull, 1910) but experienced rapid declines in subsequent decades, likely due to competition from *Z. lateralis* and increased predation following the introduction of the black rat (*Rattus rattus*) in the 1940s. By 1962, there were only an estimated 50 individuals, and by 1978, just a handful remained, culminating in its likely extinction in the following decades (Dutson, 2013; Hume & Walters, 2012).

The possibility of hybridisation among Norfolk Island *Zosterops* species was discussed by Mees (1969) and Gill (1970), who described three historical specimens collected in 1912 and 1913, purported to be hybrids between *Z. lateralis* and *Z. tenuirostris* based on intermediate body size, colouration and wing formula. Given the rarity of intermediate specimens in the early-20th century collections and lack of intermediates found subsequently, both authors concluded that hybridisation between the two species occurred infrequently soon after secondary contact, but quickly stopped altogether. Grant (1978) also dismissed ongoing hybridisation as a mechanism to explain changing bill dimensions in Norfolk Island *Z. lateralis* over time, citing a lack of intermediate specimens. Therefore, this case of hybridisation was relegated to a curious natural history note, without further consideration of evolutionary consequences for either species. However, the phenotypic and genetic characteristics of the Norfolk Island *Z. lateralis* population suggest that introgressive hybridisation may have shaped their evolution. In addition to the morphological differences evolved over only a century (Clegg, Degnan, Moritz, et al., 2002), analysis of RADseq data showed that the Norfolk Island population had a distinct genetic profile despite being the most recent in the natural colonisation sequence (Sendell□Price et al., 2021). Genetic drift due to population founding likely contributed to this, but comparable bottlenecks (e.g., in Tahiti) have not produced the same degree of genetic distinctiveness (Sendell□Price et al., 2021). Given the existence of putative hybrid specimens from the early 20th century, introgression from an endemic *Zosterops* species remains a possibility to explain the distinctive genetic profile of the Norfolk Island *Z. lateralis* population. *Z. albogularis* was not implicated in the hybridisation narrative outlined by Gill (1970), however, its plumage is sufficiently similar to that of *Z. lateralis* that, despite their size difference, misidentifications have been highlighted in the literature (Mees 1969). Therefore, the potential for introgressive hybridisation among the three Norfolk Island *Zosterops* species raises questions about the maintenance of species boundaries and consequences of introgression for single island endemics, along with the possibility of incipient ‘ghost introgression’ - where extinct species are represented in living genomes (Ottenburghs, 2020). Here, we utilise the Norfolk Island *Zosterops* system to investigate hybridisation after a historically recent secondary sympatry event resulting from long-distance island colonisation. We generate whole genome sequences from field-collected blood samples as well as historical specimens from the early 20th century, investigate population structure, and apply introgression analyses to determine: i) the identities of putative hybrids and other historical specimens before and after *Z. lateralis* arrival; and ii) whether hybridisation resulted in backcrossing and introgression.

This work aims to inform our understanding of hybridisation dynamics in island birds and their implications for the conservation of single-island endemics.

## Materials and Methods

### Sample collection and sequencing

#### Museum samples

DNA was extracted from the toe pads of 16 historical museum specimens held at the American Museum of Natural History and the 10 samples with the highest DNA quality were sent for sequencing (Table S1). A phenol-chloroform extraction protocol was used to maximize DNA yield (Billerman & Walsh, 2019; Tsai et al., 2020). A volume of 25 μl of extracted DNA per sample was sent to Arbor Biosciences (Ann Arbor, Michigan, USA) for library preparation and sequencing using the myReads service. Libraries were prepared using a combination of single-stranded ancient DNA protocols with double-stranded genomic DNA methods. Libraries were sequenced to a target coverage of 20X on a NovaSeq 6000 S4 flow cell with 150bp paired-end reads.

### Field sampling

*Z. lateralis* (15 individuals) and *Z. tenuirostris* (7 individuals) were sampled from Norfolk Island in 1998 and the North Island of New Zealand (*Z. lateralis* only, 12 individuals) in 1997 (Table S1). Birds were caught using mist nets or traps, and 20 to 40 μl of blood collected from the brachial wing vein was stored in 1 ml of lysis buffer (Seutin et al., 1991). All sampling was non-destructive, and birds were released at the point of capture. A suite of morphological measurements was also taken from field-collected individuals: wing chord, tail, tarsus, bill length to posterior nostril opening, bill width and depth at anterior nostril opening, and weight (all measurements taken by SMC). All measurements showed a clear separation between the two species (Fig. S1, t-test p < 0.001 for all comparisons).

DNA was extracted from blood samples using standard phenol-chloroform extraction as described in Estandía et al. (2025). Library preparation and whole genome sequencing were conducted by Novogene UK and the resulting libraries were sequenced to a target coverage of 5X on the Illumina Novaseq 6000 platform (Illumina, San Diego) using paired-end 150bp sequencing reads. We downloaded publicly available reads from eight other *Zosterops* species to use as outgroups for selected analyses (Table S1).

### Cleaning of sequencing reads

#### Museum specimen reads

Museum sequencing reads were cleaned with the software nf-polish (Irestedt et al 2022, available on GitHub at https://github.com/MozesBlom/nf-polish). nf-polish is a reproducible pipeline built in Nextflow v22.04, specialised for cleaning high-throughput sequencing reads extracted from historical (museum) specimens. As part of this pipeline, data underwent adapter trimming using Trimmomatic v0.39, read merging using PEAR v0.9.11, quality trimming using Trimmomatic v0.39, and removal of low complexity reads using a custom Python script (available at https://github.com/MozesBlom/nf-polish/blob/main/bin/remove_low_complex.py). Quality checks were performed using FastQC v0.11.8 before and after cleaning (Reports available on Dryad). We chose not to carry out deduplication at this stage to avoid unnecessary discarding of reads (following Irestedt et al., 2022).

### Blood sample reads

We checked the quality of the blood sample reads using FastQC (Reports available on Dryad) and verified that there was no adapter contamination, but did not conduct other quality filtering at this stage. This avoided the unnecessary discarding of reads from our low-depth (∼4x) sequencing data.

### Outgroup and ancestral sequence

For the outgroup taxon required for introgression analyses, we downloaded raw Illumina reads from a male *Zosterops borbonicus*. We also downloaded raw Illumina reads from seven other *Zosterops* species to construct an ancestral sequence for SFS polarisation (used to calculate F_ST_). All SRA accession numbers are available in Table S1. The quality of downloaded reads was assessed from FastQC reports (Reports available on Dryad). We then trimmed reads and removed adapters using fastp v0.24.0, setting a minimum length of 30 bp and a minimum PHRED quality score of 20. We also cut the front and tail ends using a 4-bp sliding window, such that any window from either end with a mean quality score below 20 would be removed.

#### Mapping reads to the reference genome

We mapped both historical and blood sample reads to the chromosome-level assembly for *Zosterops lateralis,* which contains both the Z and W chromosomes (Clegg et al., 2025). For mapping, we used a reproducible mapping pipeline built in Snakemake v8.28.0 (available on GitHub at https://github.com/abbyevewilliams/map-pipeline). Briefly, the pipeline maps reads to the reference genome using BWA-mem2 v2.2.1, sorts and indexes the reads using Samtools v1.21, marks duplicates using PicardMarkDuplicates v3.3.0 (or DeDup v0.12.8 for historical samples), and calculates summary statistics using Samtools v1.21. Finally, for historical samples, we assessed age-related damage using MapDamage2 v2.2.2 and produced bam files with rescaled quality scores for further analysis.

#### Variant calling

To call variants on the aligned BAM files, we used PopGLen v0.4.1 (Nolen, 2025), a reproducible pipeline specialised for calling variants within a genotype likelihood (GL) framework that accounts for genotype uncertainty in low-depth sequencing data. The configuration file containing all settings used is available with the stored code (see Data Accessibility and Benefit-Sharing). Briefly, we removed reads mapping to unplaced scaffolds (including the mitochondrion) and to regions with extremely high or low depth (the top/bottom 1% of depth, calculated separately for blood samples vs historical samples). We also filtered out reads with mapping or base quality scores < 20. Because the blood and historical samples had different average depths after filtering (blood: 3.8X, historical: 14.2X, Fig. S2), we downsampled the historical samples to an average depth of 4X to avoid biasing the variant calling procedure towards higher depth samples.

Given the low depth of our sequence data, we did not call genotypes directly; instead, we used ANGSD v0.940 (Korneliussen et al., 2014), invoked by PopGLen, to call GLs for each dataset, using the parameters specified in the PopGLen configuration file. As most programs do not take GLs as input, we then generated a ‘hard-call’ dataset for downstream analyses. We used a custom Python script written by Andrea Estandía (available at https://github.com/abbyevewilliams/norfolk-hybrids/blob/main/hard_call/beagle2VCF.py) to convert the ANGSD-outputted beagle file to a VCF, whilst retaining only those sites where genotype probabilities for the most likely genotype were ≥ 0.8 for at least 90% of individuals.

The most likely genotypes were taken as hard-called genotypes for downstream analyses. This dataset contained 1,673,542 SNPs, which we filtered as per the requirements of each downstream analysis (Table S2). Where possible, we compared the results of analyses using the hard-call dataset with those using the GL dataset.

#### Relatedness and population structure

Using PLINK v1.9 (www.cog-genomics.org/plink/1.9; Chang et al., 2015), we filtered the hard-call dataset for population-structure analyses. We removed sex chromosomes, then pruned for linkage disequilibrium (LD), removing SNPs with r2> 0.2 in 50kb windows. Both of these filtering steps are important to ensure independence among loci, which is violated in sex chromosomes due to their distinct inheritance patterns and demographic histories, as well as in regions of linkage disequilibrium. We also pruned for Hardy-Weinberg disequilibrium in PLINK v2.0, selecting a p-value of 10□□ and k=0.001, such that SNPs were removed if their exact Hardy-Weinberg p-values fell below p·10 nk. We kept SNPs with a lack of heterozygotes as these may be indicative of population structure, and employed the ‘midp’ modifier, which reduces the tendency of the filter to favour sites with missing data (Graffelman & Moreno, 2013). We pruned separately for the full dataset (all individuals) and for a Norfolk Island-only dataset, leaving 371,918 unlinked SNPs in the full dataset and 224,573 in the Norfolk Island-only dataset.

Using the hard-call, filtered dataset containing all individuals, we proceeded with population structure analyses. We first calculated KING-relatedness (Manichaikul et al., 2010) for all sample pairs. Two individuals (AEB01_L0003 and AEB01_L0004) had a kinship coefficient of 0.11, consistent with a second-degree relationship (e.g., half siblings), and were therefore treated as related in later analyses. We performed a Principal Component Analysis (PCA) in PLINK, decomposed the covariance matrix into eigenvectors, and plotted the two eigenvectors that explained the most variation. We also performed an ADMIXTURE (Alexander et al., 2009) analysis with 10-fold cross-validation for k=1 to k=10, then visualised log-likelihood increase as an elbow plot, selecting the value of k at which log likelihood ceased to decrease sharply. These analyses were performed with and without AEB01_L0004, and neither condition affected the results (Fig. S3). We used the packages *tidyverse* and *ggplot2* in R v4.1 to plot results from this and all other analyses.

To verify that our population structure results were replicable in a GL framework as well as from hard-calls, we repeated the population structure analyses in ANGSD. We first pruned the ANGSD-outputted beagle files for linkage disequilibrium using ngsLD (Fox et al., 2019), leaving 538,145 sites. We then ran PCAngsd (Meisner & Albrechtsen, 2018) for PCA and NGSadmix (Skotte et al., 2013) for clustering on the Beagle file of GLs. Population structure results were similar when produced with this GL framework (Fig. S4).

#### Genetic divergence

To quantify divergence between *Z. lateralis* and *Z. tenuirostris*, we calculated F_ST_ using a site frequency spectrum (SFS)-based approach implemented in ANGSD, inferring allele frequency likelihoods from GLs to account for uncertainty in genotype estimation. We compared the 1990s *Z. tenuirostris* with Norfolk Island *Z. lateralis* but did not involve *Z. albogularis* in any comparisons due to a small sample size (n=2). We were unable to calculate other measures of population divergence such as Nei’s (1987) D_xy,_ as this relies on invariant sites which cannot be confidently detected with low-depth data.

To calculate F_ST_, we first calculated the site allele frequencies (SAF) for both species, using a mapping and base quality threshold of 30, and removing reads with multiple best hits or no correctly mapped mate. We also removed triallelic sites and sites with depth <10 or >100 across all samples. We additionally set the minimum number of individuals at a site to 7 for *Z. lateralis* and 4 for *Z. tenuirostris*. The SAF was polarized based on an ancestral sequence generated from eight non-Australian *Zosterops* species (Table S1) using ANGSD’s doFasta command. We set a minimum mapping and base quality of 25 and removed reads with multiple best hits or with an unmapped mate. We used the SAF to calculate the 2D SFS, and used ANGSD’s realSFS command to obtain a genome-wide estimate of F_ST_ which is robust to genotype uncertainty in low-depth sequencing data.

#### Mitochondrial haplotype network

To create a full mitochondrial alignment, we used ANGSD to extract mitochondrial reads from the alignment and create a FASTA file used as input for the alignment software mafft v7.725 (Katoh & Standley, 2013). Relationships among mitochondrial sequences were visualised as a mitochondrial haplotype network with PopArt v1.7 (Leigh & Bryant, 2015) using a median neighbour-joining tree. We set the epsilon parameter to 0, such that only the shortest, most parsimonious connections are retained and the resulting network is as tree-like as possible.

#### Phylogenomic analysis

From the hard-call dataset, we constructed a maximum-likelihood tree using IQ-TREE v2.4.0 (Minh et al., 2020). To prepare the data, we pruned for LD using PLINK v1.9 and thinned to 100,000 SNPs. We converted our VCF file to the PHYLIP format using the vcf2phylip.py script (https://github.com/edgardomortiz/vcf2phylip). We ran IQTREE, selecting the best model using ModelFinder (Kalyaanamoorthy et al., 2017). As SNP data do not contain invariant sites, we employed the +ASC option to account for ascertainment bias. We also assessed node support using an implementation of the SH-aLRT test (Guindon et al., 2010) with 1000 replicates, as well as 1000 ultrafast bootstraps, adding the -bnni flag to avoid artificial inflation of bootstrap support values (Hoang et al., 2018). Trees were visualised using the R packages *ggtree* v4.1.1 (Yu et al., 2017) and *ape* v5.8-1 (Paradis & Schliep, 2019).

#### Sex determination

In birds, females are the heterogametic sex (i.e., WZ are females; males are ZZ) (Takagi & Sasaki, 1974). To determine sex from sequencing reads, we calculated the ratio of the number of W chromosome reads to Z chromosome reads. Individuals with a ratio of ∼0 were identified as male, and individuals with a ratio of >>0 were identified as female (following Kersten et al., 2023, Table S3). This was carried out on the hard-call dataset.

#### Hybridisation analyses

To verify hybrid identities for genotypically intermediate individuals, we ran two programs on the hard-called, LD- and HW-pruned dataset (Norfolk Island individuals only): HyDe (Blischak et al., 2018), and triangulaR (Wiens et al., 2025). HyDe tests for hybridisation using quartet-based ‘phylogenetic invariants’, focusing on SNP site patterns whose expected frequencies are identical across all alternative quartet topologies under the null model, and deviations from these expectations are used to detect hybrid ancestry. TriangulaR uses ‘ancestry-informative markers,’ sites differing strongly between populations, to calculate the relationship between hybrid index, the estimated proportion of an individual’s genome inherited from one species, and interclass heterozygosity, the heterozygosity resulting from alleles from two different species. The relationship between these two variables can be used to assign hybrid individuals to a class (e.g. F1, F2, backcross). We ran TriangulaR using an allele frequency threshold of 0.9.

#### Introgression analyses

We used the Dsuite package (Malinsky et al., 2021) to test for signals of introgression between the *Zosterops* species on Norfolk Island. We used the hard-call dataset to calculate the D-statistic (Durand et al., 2011; Green et al., 2010) and admixture proportion (or *f_4_*-ratio) (Patterson et al., 2012). We used the endemic species (*Z. tenuirostris/Z. albogularis*) as the ‘donor’ population (P3), Norfolk Island Z. lateralis as the ‘receiver’ population (P2), New Zealand *Z. lateralis* as P1 (the ‘control’/non-admixed population), and *Z. borbonicus* as the outgroup (though we tested all possible combinations; results shown in Table S4). For *Z. tenuirostris*, we ran Dsuite separately using the historical (1906/1926) and blood sample (1998) reads for this species, but achieved similar results (Table S4).

For significant Dsuite comparisons, the genomics_general library (https://github.com/simonhmartin/genomics_general) was used to calculate ABBA-BABA statistics (D, f_D_) in 50kb sliding windows with a 10kb step size across the genome, using only windows with at least 50 SNPs. We focus on f_D_ results for sliding-window comparisons because D is biased in small genomic regions (Martin et al., 2015). As f_D_ is not valid for windows where D is negative, these windows were excluded from further analyses. To complement and verify our sliding window analysis, we also used Twisst2 (Martin, 2026), a topology-weighting software that calculates support for different genealogical relationships across the genome.

Patterson’s *D*-statistic (including f_D_, f_DM_) does not provide information regarding the timing of introgression. We therefore utilised a simple extension of the *D*-statistic, the *D* frequency spectrum (*D*_FS_) developed by Martin & Amos (2021), in which *D* was partitioned according to the frequencies of derived alleles. Like the site frequency spectrum (SFS), the *D*_FS_ is sensitive to allele frequencies: recent or ongoing introgression produces a peak in *D*_FS_ among low-frequency alleles, whereas past introgression produces a peak in *D*_FS_ among high-frequency alleles as introgressed alleles are either driven towards fixation by drift or selection or are lost in the recipient population. Peaks at high-frequency alleles can also be indicative of a recent bottleneck, known to have occurred in Norfolk Island *Z. lateralis* (Martin & Jiggins, 2017). The *D*_FS_ was calculated using additional Python scripts developed by S. Martin (https://github.com/simonhmartin/dfs). We only calculated the *D*_FS_ for the *Z. lateralis* x *Z. tenuirostris* comparison, owing to the small sample size (n=2) of *Z. albogularis*.

We did not perform introgression analyses that required haplotype-phased genomes (such as local ancestry inference (e.g., RFMix2, Lisi & Campbell, 2024) and chromosome painting (e.g., ChromoPainter, Lawson et al., 2012)) as our genomes were not phased and we could not reliably perform statistical phasing (e.g., ShapeIT, Delaneau et al., 2012) given the sequencing depth of the data.

#### Functional analysis of introgressed regions

To explore potential functional roles of introgressed regions, we identified genes falling within the highest-scoring f_D_ windows. We selected the top 1% of f_D_ windows for each species comparison, created a FASTA file for these sequences (masked for repetitive regions), and used BLAST (Altschul et al., 1990) to search them against the annotated *Taeniopygia guttata* reference genome. We filtered hits to those with an e-value below 1e-6, with alignments at least 100 bp long and at least 60% similarity. We tested functional roles of gene candidates using two complementary methods. First, we ran a gene set enrichment analysis using g:Profiler (Raudvere et al., 2019), which tests whether particular biological functions are over-represented in a given gene set. Second, we checked for overlap with GePheBase (Courtier-Orgogozo et al., 2020), a database of gene-phenotype associations, using bird entries as candidates.

## Results

### Population structure

Genome-wide PCA on the Norfolk Island samples revealed three distinct genetic clusters reflecting the three species: *Z. lateralis*, *Z. tenuirostris* and *Z. albogularis* (Fig. 2A). We did not observe substructure based on specimen age (blood samples vs historical specimens). The three putative *Z. lateralis x Z. tenuirostris* phenotypic hybrids recovered in an intermediate position between the *Z. lateralis* and *Z. tenuirostris* clusters. Unexpectedly, one additional historical specimen recovered in an intermediate position: AEB01_L0007, collected in 1926 and identified as *Z. lateralis* in AMNH specimen labels, was intermediate between *Z. lateralis* and *Z. albogularis* in the PCA. The three species groupings were corroborated by admixture analysis: clustering with ADMIXTURE revealed k=3 as the optimal number of clusters based on log-likelihood increase (Table S5), and these clusters corresponded to each species group, as well as highlighting the four individuals with intermediate PCA position as mixed-ancestry individuals (Fig. 2B). The mitochondrial haplotype network also revealed three major nodes (Fig. 2C). All four putative hybrids were clustered around or within the *Z. lateralis* node, suggesting *Z. lateralis* maternal identity.

**Figure 2.**
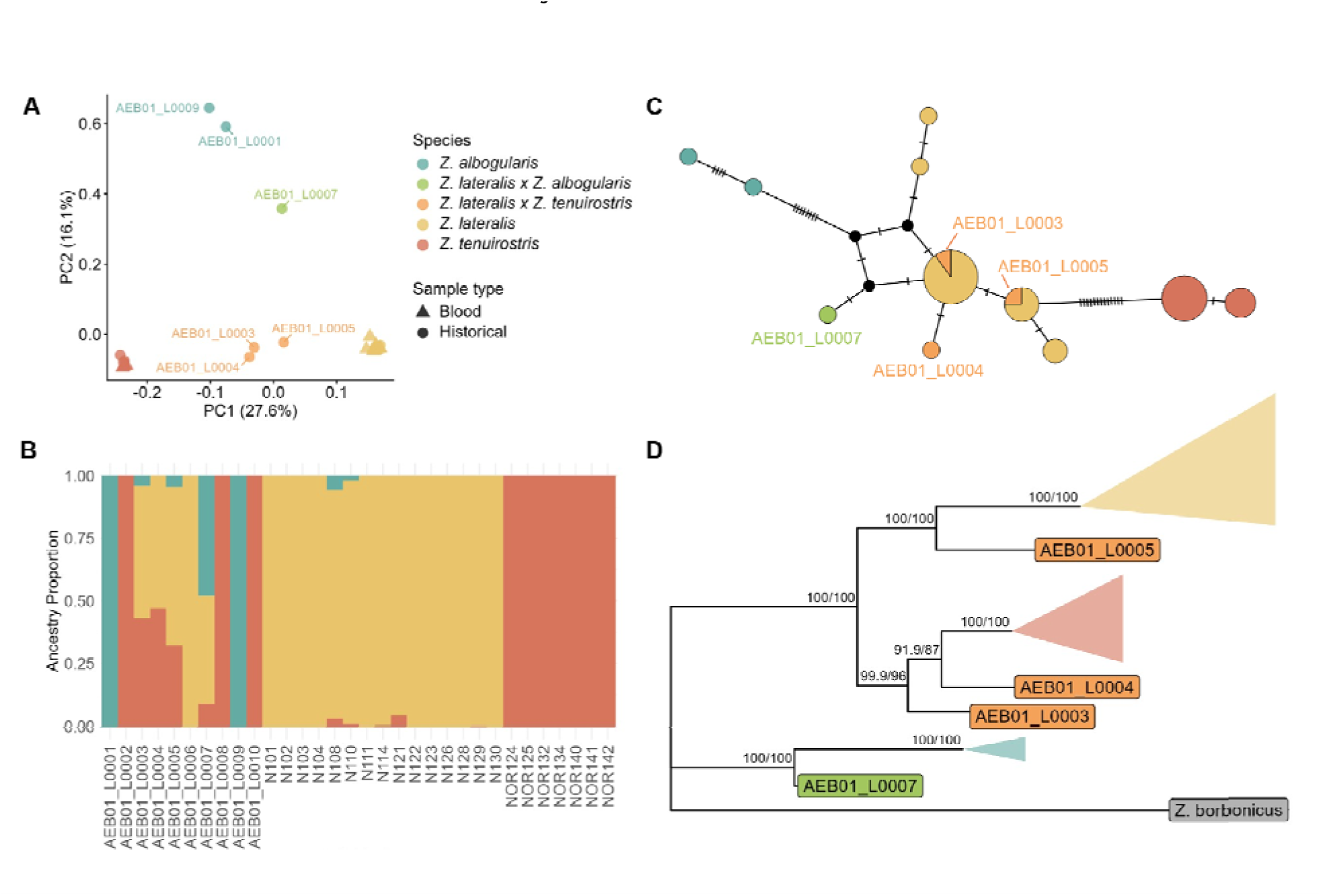
A) PCA based on 224,575 unlinked SNPs showed three main clusters reflecting the species groups (*Z. albogularis*: teal, *Z. lateralis*: yellow, *Z. tenuirostris*: red) as well as individuals with mixed ancestry (*Z. lateralis x Z. albogularis;* light green, *Z. lateralis x Z. tenuirostris*, orange). Blood samples are indicated by triangles and historical samples with circles. Related individuals are included. B) Admixture plot with k=3 showing three main groupings with sample names shown. C) Median joining mitochondrial haplotype network constructed in PopArt based on mitochondrial alignment. Dashes represent nucleotide differences between clusters. Hybrid samples are labelled. D) Maximum likelihood tree constructed in IQTREE based on 100,000 unlinked autosomal SNPs with bootstrap support (SH-aLRT/UFboot) shown for main nodes. Main species groupings are collapsed for clarity. Outgroup (*Z. borbonicus*) is shown in grey.

Finally, the genome-wide maximum likelihood phylogeny supported three major clades corresponding to each species, with putative hybrids positioned as outgroups to these clades (Fig. 2D). All major nodes were strongly supported (ultrafast bootstrap ≥91% and SH-aLRT ≥87%). The divergence between *Z. lateralis* and *Z. tenuirostris* was supported by a genome-wide F_ST_ of 0.41, indicating moderate-high divergence, though this may have been inflated by the sequential genetic bottlenecks experienced by *Z. lateralis*.

### Sex determination

Based on the ratio of reads mapping to the W and Z chromosomes, we could confidently determine the sex of 8/10 historical and 22/22 blood samples (Table S3). Of the historical samples that contained putative hybrids, 6 were male, 2 were female, and 2 were ambiguous (likely representing females whose W chromosomes were degraded due to age; all collected in 1926). All 3 of the *Z. lateralis x Z. tenuirostris* putative hybrids were males, whereas the *Z. lateralis x Z. albogularis* putative hybrid was ambiguous.

### Hybridisation

Hybridisation detection with HyDe confirmed hybrid identity for the 4 genomically intermediate individuals (Table 1). Triangle plots based on 4,270 sites for the *Z. lateralis x Z. tenuirostris* comparison and 8,389 sites for the *Z. lateralis x Z. albogularis* comparison shows all hybrids were early-generation (F1- or F2-like, Fig. 3).

**Figure 3.**
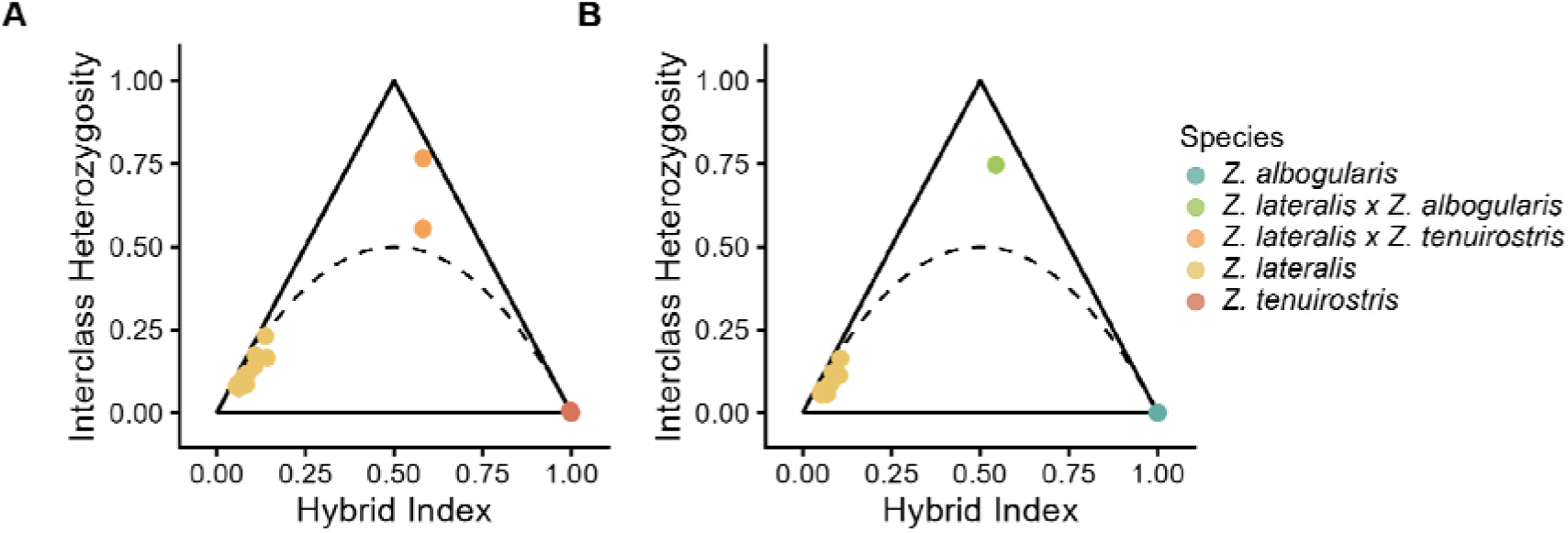
A) Triangle plots for the *Z. lateralis x Z. tenuirostris* comparison based on 4,270 ancestry-informative markers (AIMs), with one related individual (AEB01_L0004) removed. B) *Z. lateralis-Z. albogularis* comparison based on 8,389 AIMs.

**Table 1.** Hybridisation detection results from HyDe. Only individuals with significant p-values are shown, corresponding to the four hybrids. The Gamma parameter describes the proportion of the genome estimated to be inherited from either parent population: a value closer to 0 indicates more P1 identity, whereas closer to 1 indicates more P2 identity. *p-values outputted by HyDe were infinitesimal for all hybrid individuals.

| P1 | Sample ID | P2 | Zscore | P-value | Gamma |
| --- | --- | --- | --- | --- | --- |
| *Z. lateralis*<br>(Norfolk<br>Island) | AEB01_L000<br>3 | *Z.*<br>*tenuirostris* | 29.0 | 0* | 0.355 |
| *Z. lateralis*<br>(Norfolk<br>Island) | AEB01_L000<br>4 | *Z.*<br>*tenuirostris* | 29.7 | 0 | 0.345 |
| *Z. lateralis*<br>(Norfolk<br>Island) | AEB01_L000<br>5 | *Z.*<br>*tenuirostris* | 23.5 | 0 | 0.424 |
| *Z. lateralis*<br>(Norfolk<br>Island) | AEB01_L000<br>7 | *Z.*<br>*albogularis* | 31.4 | 0 | 0.352 |

### Introgression

Dsuite analysis indicated low but significant levels of introgression into *Z. lateralis* from both endemic species, *Z. tenuirostris* and *Z. albogularis*. We recovered a signal of introgression when using Norfolk Island *Z. lateralis* as P2 (the receiver population), and historical (i.e. non-admixed) *Z. tenuirostris* or *Z. albogularis* as P3 (the donor population; Fig. 4A). For both *Z. tenuirostris* and *Z. albogularis*, there was a higher than expected number of alleles shared with *Z. lateralis* than would be expected under a neutral scenario, resulting in D statistic that was significantly greater than zero (Fig. 4B; for *Z. lateralis x Z. tenuirostris*: D = 0.026, P < 0.001; for *Z. lateralis x Z. albogularis*: D = 0.0016, P < 0.001). The estimated admixture fraction of the Norfolk Island *Z. lateralis* genome was estimated at 1.3% for both *Z. tenuirostris* and *Z. albogularis* contributions. However, this signal of introgression was not observed when P2 and P3 were reversed, indicating unidirectional introgression from each endemic into *Z. lateralis*. The D_FS_ (Fig. 4C) based on the *Z. lateralis x Z. tenuirostris* comparison supported recent introgression, with a peak at low-frequency alleles, consistent with the historically recorded timing of contact between the species. The D_FS_ also showed a smaller peak at high-frequency alleles, consistent with a population bottleneck prior to gene flow. The distribution of sliding-window f_D_ scores indicated low but consistent introgression across the entire genome (Fig. 4D), a result corroborated by the topology weighting analysis (Fig. S5).

**Figure 4.**
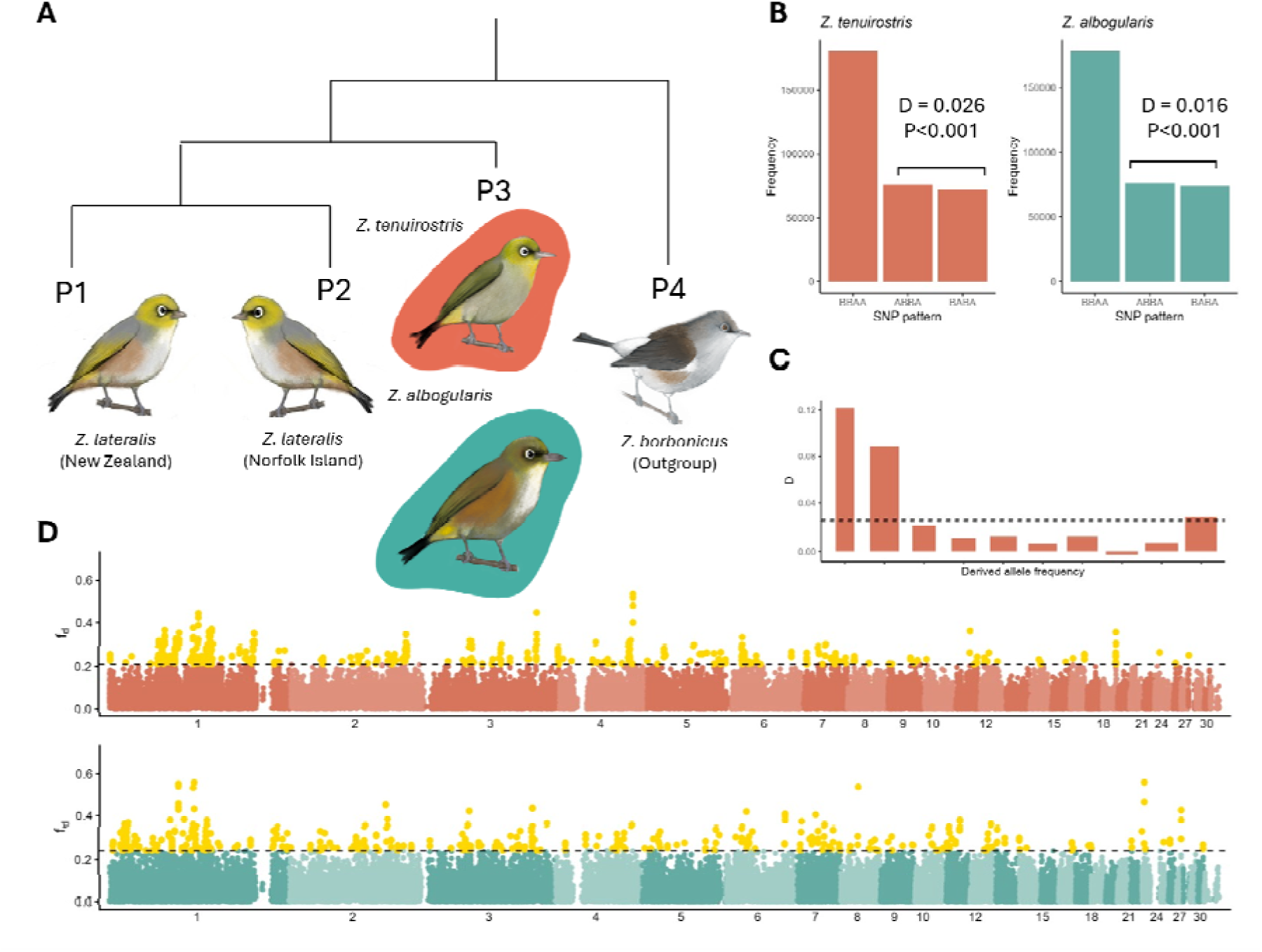
A) Topologies tested for Dsuite Dtrios (P1 = *Z. lateralis* (New Zealand), P2 = *Z. lateralis* (Norfolk Island), P3 = *Z. tenuirostris* or *Z. albogularis*, P4 = *Z. borbonicus* (outgroup). B) Numbers of BBAA, ABBA, and BABA sites in each comparison. The excess of ABBA indicates gene flow between P2 and P3 (rather than P1 and P3, which would be indicated by an excess of BABA). C) *D* frequency spectrum (*D*_FS_) for *Z. lateralis x Z. tenuirostris* comparison. Each vertical line indicates the stratified *D*-value for each derived allele frequency bin. The widths of vertical lines are proportional to their weighting. The dotted line shows the average D-value. D) f_D_ calculated in 50 kb windows across the genome (sex chromosomes omitted due to low SNP density). All yellow points above the grey dotted line are in the top 1% of outliers. Bird illustrations by AW.

### Functional analysis of introgressed regions

Filtering to the top 1% of f_D_ outliers, we recovered 401 windows for the *Z. lateralis x Z. tenuirostris* comparison and 391 windows for the *Z. lateralis x Z. albogularis* comparison. From these windows, we recovered 243 high-confidence genes for the *Z. lateralis x Z. tenuirostris* comparison and 266 for the *Z. lateralis x Z. albogularis* comparison. None of these genes were found in the GePheBase search. However, we noted that the *Z. lateralis x Z. tenuirostris* gene set contained *LEMD3*, linked to dwarfism in chickens (Perini et al., 2023), and that the highest-f_D_ window contained *RELN*, a known target of the *FOXP* pathway linked to song regulation in zebra finches (Lovell et al., 2018) and putatively linked to body size in a recent GWAS on body size on *Z. lateralis* (Estandía et al., 2023).

The g:Profiler gene set enrichment analysis, when applied to the *Z. lateralis* x *Z. tenuirostris* comparison, yielded one significant Gene Ontology category: proteasomal protein catabolic process (GO term GO:0010498, p < 0.05). This category comprised 14 genes linked to various functions, ranging from iron metabolism (*FBXL5*; Yin et al., 2019) to immunity (*TRIM13*; Zhou et al., 2024). The same analysis for the *Z. lateralis x Z. albogularis* comparison also gave one significant category, negative regulation of retinoic acid receptor signaling (GO term GO:0048387 P < 0.05). This category was linked to the genes *CNOT1* (associated with spatial cognition in chickadees, Semenov et al., 2024) and ZNF536 (associated with cold adaptation in chickens, Xu et al., 2021).

## Discussion

Whole-genome sequencing of field-collected and historical samples from a trio of *Zosterops* species on Norfolk Island has revealed the extent of introgression following a historically recorded secondary sympatry event in the early 20th century. Hybridisation between the recently colonised *Z. lateralis* population and two endemic *Zosterops* species resulted in introgression into the coloniser. This included incipient ‘ghost introgression’ with DNA from the extinct endemic, *Z. albogularis*, represented in *Z. lateralis* genomes from late 20th-century samples. Introgression was genome-wide, and functional analysis suggested a range of potential adaptive roles. The features of genomically F1-like hybrids provide clues as to how both genomic incompatibility and behavioural preferences shape introgression patterns in *Zosterops*, providing novel, potentially adaptive genes to the coloniser while maintaining the genomic integrity of the extant, narrow range endemic.

Our genomic results support a more complex scenario of hybridisation and introgression among *Zosterops* species on Norfolk Island than previously proposed. After arrival, *Z. lateralis* hybridised with both endemics, as evidenced by confirmation of genomically intermediate hybrids with *Z. tenuirostris*, collected in 1912 from at least two unrelated clutches (as shown by relatedness results from hybrids specimens), and the unexpected finding of a genomic hybrid with *Z. albogularis* from a specimen collected in 1926. Detection of genomically widespread introgressed regions of the endemic species genomes in silvereye samples collected in the late 20th century indicated that hybridisation either occurred frequently in early decades, then potentially ceased, or has continued at a low level over the century. We could not distinguish between these possibilities in this study due to the difficulty of conducting demographic analyses on short time scales with small sample sizes and relatively low sequencing depth, as well as the fragmented nature of research and monitoring of these species on Norfolk Island. However, our finding that an average of 1.3% of the genome of Norfolk Island *Z. lateralis* collected in the 1990s was derived from each of the endemic species suggests that hybridisation may have continued beyond the early 20th century, as ancestry from a single pulse of hybridisation is expected to halve each generation under random mating, leading to rapid decay of genomic contribution over time (Pool & Nielsen, 2009). Additionally, anecdotal evidence of a phenotypic hybrid caught in recent years suggests that hybridisation may have continued in recent decades (C. Young, person comm).

Introgression appears to be unidirectional from both endemics into *Z. lateralis*, suggesting preferential backcrossing of hybrids with silvereyes rather than with either endemic species. This asymmetrical pattern has several possible explanations, for example, a biased sex ratio in hybrids may have contributed to asymmetrical mating. Our genetic sexing indicates that all four hybrid samples were males, consistent with Haldane’s (1922) Rule that hybrid offspring are more likely to be viable if they are the homogametic sex (males in birds). Female *Z. lateralis* may have been more likely to accept these hybrid males as mates than females of either endemic species, for example, because in the early stages of *Z. lateralis* colonisation, low *Z. lateralis* numbers may have promoted acceptance of heterospecific partners, including hybrids. Wirtz (1999) proposes that such an imbalance in numbers could lead to unidirectional hybridisation as a general outcome of sympatry. However, as *Z. lateralis* was already more common than either endemic species by 1910 (Mees, 1969), this ‘desperation hypothesis’ (Hubbs, 1955) would have applied only for a short time. Hybrids may also have been more attractive to female *Z. lateralis* because of their larger size (Grant & Grant, 1997) or other intermediate characteristics such as plumage (Gill, 1970). Large body and bill size have been shown to be favoured by natural selection in multiple island populations of silvereyes (Clegg, Degnan, Kikkawa, et al., 2002; Clegg et al., 2008) and are consistent with the direction of evolution expected under the island rule (Jezierski et al., 2024).

The direction of hybrid backcrossing may also have been influenced by the parental identities of the hybrids. Our mitochondrial haplotype network suggested that all four hybrid specimens had *Z. lateralis* maternal ancestry, which may have implications for hybrid mate choice preferences. Grant and Grant’s (1997) work on *Geospiza* finches shows that courtship behaviour can result from parental imprinting, for example in *G. fortis*, females may pair with heterospecific males singing a similar song to their fathers, or with similar morphology to their mothers. Though parental imprinting has not, to our knowledge, been studied in *Zosterops*, male hybrid offspring imprinting on their *Z. lateralis* mother’s characteristics could lead them to seek *Z. lateralis* mates. However, our hybrid sample size is small, and verifying our interpretation would require more sequencing of possible hybrids. Analysis of traits involved in maintaining species boundaries (e.g., song, courtship behaviour or plumage) would further clarify the mechanisms underlying the directionality of backcrossing.

The introgression of *Z. albogularis* genetic variation into *Z. lateralis* provides an uncommon example of recently acquired ‘ghost introgression’ (Ottenburghs, 2020). Cited examples of ghost introgression have occurred over more ancient time scales, such as the hybridisation of humans with Neanderthals (> 30,000 years ago; Green et al., 2010), brown bears with extinct cave bears (> 25,000 years ago; Barlow et al., 2018), and bonobos with an extinct ape lineage (377-637 thousand years ago; Kuhlwilm et al., 2019). To our knowledge, ghost introgression has not previously been demonstrated to occur over hundreds of years, but future sequencing of more recently extinct species from museum specimens may reveal that it is a more widespread phenomenon. Although *Z. albogularis* is already likely extinct, with no verified sightings since 1979 (Schodde et al., 1983), reconstructing the genomic circumstances surrounding its extinction may still provide useful insights for the conservation of extant species. Our introgression analyses did not detect gene flow from *Z. lateralis* to *Z. albogularis*, though our data do not capture the mid-20th century, when this species was rapidly declining, and mate choice options may have been restricted. Sequencing of more recent *Z. albogularis* specimens could clarify if this direction of introgression occurred as the species declined toward extinction.

Introgression may play a constructive role in evolution where introgressed genes are adaptive (Jones et al., 2018; Kendall et al., 2025; Meier et al., 2017; Pardo-Diaz et al., 2012; Valencia-Montoya et al., 2020). While the direct effects of introgressed genes on fitness proxies, such as survival and reproduction, in *Z. lateralis* are outside the scope of this study, our functional analysis revealed that regions of the genome introgressed from *Z. tenuirostris* into *Z. lateralis* were enriched for genes associated with a wide range of potential functions. This contrasts with some systems in which introgression has been limited to one or a small number of specific loci with known phenotypic effects (Jones et al., 2018; Pardo-Diaz et al., 2012; Valencia-Montoya et al., 2020). However, other studies have reported similarly widespread functional roles of introgressed loci; for example, one study on introgression in big cats showed that putatively introgressed loci were enriched for functions such as craniofacial and limb development, pigmentation, reproduction, and sensory perception (Figueiró et al., 2017). Notably, in this study, we did not identify any gene candidates previously linked to bill shape, such as ALX1 (Lamichhaney et al., 2015) or BMP4 (Abzhanov et al., 2004), a feature which has changed over time in Norfolk Island *Z. lateralis* and is suggested to have resulted from early hybridisation (Grant, 1978). Our observed lack of strong, single-locus introgression peaks may be due to the relatively short timescale of secondary contact, weak selection on introgressed regions, or an artifact of relatively low sequencing depth and small sample size, which could be improved upon in future studies.

Our discovery of introgression among *Zosterops* species on Norfolk Island raises the possibility that it could be more frequent than previously supposed in groups, such as *Zosterops*, that are not known for hybridisation (Ottenburghs, 2019). Estandía et al. (2025) suggested that introgression could explain uncertainties in phylogenetic relationships among *Zosterops* species and that numerous cases of secondary sympatry warrant investigation through a genomic lens. More broadly, our findings reflect a growing body of evidence from diverse taxa where hybridisation is only detected after the application of genomic data, for example, between morphologically similar but genetically divergent lineages of American crow (Slager et al., 2020). Outside of birds, genomic data has revealed cryptic hybridisation between the endangered Booroolong frog *Litoria booroolongensis* and the eastern stony creek frog *L. wilcoxii,* which may threaten the existence of the endangered species (Liu & Rowley, 2025). These studies add to the growing body of evidence, illuminated by WGS, that hybridisation in nature has been far more common than previously supposed, and sequencing data from a wider range of taxa will provide a more accurate estimate of its true prevalence across the tree of life.

The degree and direction of hybridisation has implications for the conservation of white-eyes on Norfolk Island. As *Z. tenuirostris* is vulnerable and declining (IUCN, 2025), it is reassuring that our results indicate it is not threatened by hybridisation. Rather, the maintenance of clear morphometric phenotypes of both *Z. tenuirostris* and *Z. lateralis* at least up until the 1990s, the sharp genetic boundaries seen in population structure analyses, and the finding that its genome remains non-admixed with *Z. lateralis*, demonstrate that hybridisation and introgression levels have been insufficient to result in a hybrid swarm scenario or extinction by hybridisation, scenarios evidenced in other bird species such as the endemic Hawaiian duck *Anas wyvilliana* (C. P. Wells et al., 2019) and Seychelles turtle dove *Nesoenas picturata aldabranus* (Skerrett et al., 2001). Yet, a growing number of studies in diverse systems, including yeast (Stelkens et al., 2014) and trout (Z. R. R. Wells et al., 2019), as well as other *Zosterops* species (Vedder et al., 2022), have highlighted how introgression can facilitate genetic rescue by providing additional genetic diversity to allow response to natural selection. Therefore, if the *Z. tenuirostris* population declines further and introgression from *Z. lateralis* occurs, it may not necessarily be detrimental.

Introgression of genetic material from both endemics, including one that is now extinct, raises an important conservation point: *Z. lateralis* on Norfolk Island is not simply another population of a widespread species, but the bearer of a unique genetic legacy shaped by historical hybridisation. Recognising such genetically important populations is crucial for conserving biodiversity at the genomic level, and differs from the ‘evolutionary significant unit’ (ESU) approach for delineating conservation units below the species level (Ryder, 1986). Although ESUs have typically been defined by reproductive isolation and genetic divergence, there is increasing recognition that no single criterion can be used to accurately decide the conservation value of a species (Fraser & Bernatchez, 2001). Norfolk Island *Z. lateralis* represents one such population which may not traditionally have been considered an ESU (it is only moderately diverged from its source population and unlikely to be reproductively isolated), but our genomic analyses have demonstrated that this population acts as a reservoir of historical biodiversity that would have been lost if not for admixture prior to the extinction of *Z. albogularis*. Such reservoirs of diversity may become increasingly important in a rapidly changing world, and must be explicitly recognised in conservation frameworks.

Several interpretations from our study were only possible in light of comparison with historical specimens from before and soon after *Z. lateralis* colonisation. Our study therefore reinforces the value of museum collections as valuable baseline sources of DNA for monitoring contemporary populations (Nakahama, 2021; Yeates et al., 2016). Furthermore, our work shows that museum collections may hold previously overlooked but important specimens, demonstrated by our discovery of an early-generation *Z. lateralis x Z. albogularis* hybrid collected in 1926 (sample no. AEB01_L0007) that had been identified as *Z. lateralis*, the true identity of which may have been obscured by the superficial phenotypic resemblance between *Z. lateralis and Z. albogularis* (Mees 1969). This particular specimen fits into a wider landscape of singular museum specimens held in collections that can provide insights into lineages no longer found in the wild, such as the Guanacaste hummingbird *Amazilia alfaroana,* which is known from a single type specimen collected in 1896 and is likely extinct (Kirwan & Collar, 2016). Museomics approaches can therefore provide exciting new insights into long-held collections and novel interpretations of the populations and species they represent.

## Acknowledgements

Whole genome sequencing of museum samples was funded by contributions from the Oxford University Press John Fell Fund, start-up funds from the Department of Zoology, Oxford, and funding from St Anne’s College, Oxford to SMC; and a British Ornithologists’ Union small grant to AE. AW is supported by the Robert and Valerie Appleby Research Scholarship. Samples from Norfolk Island were collected with permissions from Norfolk Island Parks and Forestry, Environment Australia (Norfolk Island National Park and Botanic Garden Act 1984), and the University of Queensland ethics clearances ZOO/520/96/ARC/PHD and ZOO/520/97/ARC/PHD. Samples from New Zealand were collected with permission from the Department of Conservation Te Papa Atawhai. We acknowledge the late Prof Jiro Kikkawa for assistance with Norfolk Island fieldwork and Prof Bruce Robertson and Fiona Robertson for assistance with New Zealand fieldwork. We are grateful to Paul Sweet and Thomas J. Trombone from the American Museum of Natural History (AMNH) in New York, New York, USA for providing access to historical samples.

## Data Accessibility and Benefit-Sharing

### Data Accessibility

Raw sequence reads (FASTQ files) are deposited in the SRA under BioProject IDs PRJNA1264480 (samples N114, N122, N126, N129, NOR124 and NOR132) and PRJNA1461875 (all other samples). Sample metadata can be found in the corresponding SRA BioProject.

FastQC reports and unfiltered BEAGLE/VCF files are available on Dryad.

Code for mapping reads is available at: https://github.com/abbyevewilliams/map-pipeline Code for downstream analyses is available at: https://github.com/abbyevewilliams/norfolk-hybrids

### Benefit–Sharing Statement

This work is the result of collaboration between researchers in four different countries and our main collaborators have been credited as co-authors. The results of this research have been shared with the broader scientific community as well as the general public. This work addresses an important conservation concern in a vulnerable species, with implications for conservation science more widely. Finally, all data and code have been made available for use by the scientific community.

## Author Contributions

Conceptualisation: SMC, ASP Funding acquisition: SMC, AE

Data collection (field sampling): SMC Data collection (museum samples): AE

Data collection (laboratory work): SMC, ASP, AMC Analysis: AW, ASP

Writing: AW, ASP, SMC Supervision: SMC, AE, DF, KR

Editing: all authors

## Notes

### Competing Interest Statement

The authors have declared no competing interest.

