## Supplementary Materials for "Recent hybridisation and ghost introgression among a trio of island passerines"

**Table S1:** the samples used in the analyses. *AEB01_L0007 was originally classified as *Z. lateralis* but our analyses show it is likely a first-generation *Z. lateralis x Z. albogularis* hybrid. ** DNA was extracted but not sequenced for 6 museum specimens, acknowledged here.

| **Sample** | **Species** | **Location** | **Year** | **Source** | **Voucher ID** | **SRA Accession** |
| --- | --- | --- | --- | --- | --- | --- |
| AEB01_L0001 | *Z. albogularis* | Norfolk Island | 1903 | Museum | AMNH 701357 |  |
| AEB01_L0002 | *Z. tenuirostris* | Norfolk Island | 1906 | Museum | AMNH 701317 |  |
| AEB01_L0003 | *Z. lateralis x tenuirostris* | Norfolk Island | 1912 | Museum | AMNH 701304 |  |
| AEB01_L0004 | *Z. lateralis x tenuirostris* | Norfolk Island | 1912 | Museum | AMNH 701305 |  |
| AEB01_L0005 | *Z. lateralis x tenuirostris* | Norfolk Island | 1913 | Museum | AMNH 701139 |  |
| AEB01_L0006 | *Z. lateralis lateralis* | Norfolk Island | 1926 | Museum | AMNH 212573 |  |
| AEB01_L0007 | *Z. lateralis x albogularis** | Norfolk Island | 1926 | Museum | AMNH 212574 |  |
| AEB01_L0008 | *Z. tenuirostris* | Norfolk Island | 1926 | Museum | AMNH 212653 |  |
| AEB01_L0009 | *Z. albogularis* | Norfolk Island | 1926 | Museum | AMNH 212667 |  |
| AEB01_L0010 | *Z. tenuirostris* | Norfolk Island | 1926 | Museum | AMNH  701316 |  |
| ** | *Z. albogularis* | Norfolk Island | 1906 | Museum | AMNH 701354 |  |
| ** | *Z. albogularis* | Norfolk Island | 1906 | Museum | AMNH 701358 |  |
| ** | *Z. lateralis lateralis* | Norfolk Island | 1926 | Museum | AMNH 212575 |  |
| ** | *Z. tenuirostris* | Norfolk Island | 1926 | Museum | AMNH 212652 |  |
| ** | *Z. albogularis* | Norfolk Island | 1926 | Museum | AMNH 212666 |  |
| ** | *Z. albogularis* | Norfolk Island | 1926 | Museum | AMNH 212670 |  |
| N101 | *Z. lateralis* | Norfolk Island | 1998 | Wild-caught |  |  |
| N102 | *Z. lateralis* | Norfolk Island | 1998 | Wild-caught |  |  |
| N103 | *Z. lateralis* | Norfolk Island | 1998 | Wild-caught |  |  |
| N104 | *Z. lateralis* | Norfolk Island | 1998 | Wild-caught |  |  |
| N108 | *Z. lateralis* | Norfolk Island | 1998 | Wild-caught |  |  |
| N110 | *Z. lateralis* | Norfolk Island | 1998 | Wild-caught |  |  |
| N111 | *Z. lateralis* | Norfolk Island | 1998 | Wild-caught |  |  |
| N114 | *Z. lateralis* | Norfolk Island | 1998 | Wild-caught |  | SRX28952079 |
| N121 | *Z. lateralis* | Norfolk Island | 1998 | Wild-caught |  |  |
| N122 | *Z. lateralis* | Norfolk Island | 1998 | Wild-caught |  | SRX28952080 |
| N123 | *Z. lateralis* | Norfolk Island | 1998 | Wild-caught |  |  |
| N126 | *Z. lateralis* | Norfolk Island | 1998 | Wild-caught |  | SRX28952081 |
| N128 | *Z. lateralis* | Norfolk Island | 1998 | Wild-caught |  |  |
| N129 | *Z. lateralis* | Norfolk Island | 1998 | Wild-caught |  | SRX28952082 |
| N130 | *Z. lateralis* | Norfolk Island | 1998 | Wild-caught |  |  |
| NOR124 | *Z. tenuirostris* | Norfolk Island | 1998 | Wild-caught |  | SRX28952084 |
| NOR125 | *Z. tenuirostris* | Norfolk Island | 1998 | Wild-caught |  |  |
| NOR132 | *Z. tenuirostris* | Norfolk Island | 1998 | Wild-caught |  | SRX28952085 |
| NOR134 | *Z. tenuirostris* | Norfolk Island | 1998 | Wild-caught |  |  |
| NOR140 | *Z. tenuirostris* | Norfolk Island | 1998 | Wild-caught |  |  |
| NOR141 | *Z. tenuirostris* | Norfolk Island | 1998 | Wild-caught |  |  |
| NOR142 | *Z. tenuirostris* | Norfolk Island | 1998 | Wild-caught |  |  |
| PN1 | *Z. lateralis* | New Zealand (North Island) | 1997 | Wild-caught |  | SRX28952023 |
| PN2 | *Z. lateralis* | New Zealand (North Island) | 1997 | Wild-caught |  |  |
| PN3 | *Z. lateralis* | New Zealand (North Island) | 1997 | Wild-caught |  |  |
| PN6 | *Z. lateralis* | New Zealand (North Island) | 1997 | Wild-caught |  |  |
| PN7 | *Z. lateralis* | New Zealand (North Island) | 1997 | Wild-caught |  |  |
| PN10 | *Z. lateralis* | New Zealand (North Island) | 1997 | Wild-caught |  | SRX28952024 |
| PN11 | *Z. lateralis* | New Zealand (North Island) | 1997 | Wild-caught |  | SRX28952025 |
| PN12 | *Z. lateralis* | New Zealand (North Island) | 1997 | Wild-caught |  | SRX28952026 |
| PN13 | *Z. lateralis* | New Zealand (North Island) | 1997 | Wild-caught |  |  |
| PN15 | *Z. lateralis* | New Zealand (North Island) | 1997 | Wild-caught |  |  |
| PN16 | *Z. lateralis* | New Zealand (North Island) | 1997 | Wild-caught |  |  |
| PN17 | *Z. lateralis* | New Zealand (North Island) | 1997 | Wild-caught |  |  |
|  | *Z. borbonicus* | Reunion | 2009 | Publicly available |  | SRX9098817 |
|  | *Z. hypoxanthus* | Papua New Guinea | 1994 | Publicly available |  | SRX6695485 |
|  | *Z. japonicus* | Not provided | Not provided | Publicly available |  | SRX9708198 |
|  | *Z. mauritianus* | Mauritius | Not provided | Publicly available |  | SRX9088330 |
|  | *Z. olivaceus* | Reunion | 2015 | Publicly available |  | SRX9196374 |
|  | *Z. pallidus* | South Africa | 2015 | Publicly available |  | SRX9239935 |
|  | *Z. poliogastrus* | Ethiopia | 2017 | Publicly available |  | SRX19767761 |
|  | *Z. virens* | South Africa | 2014 | Publicly available |  | SRX9202113 |

**Table S2**: the datasets used in each analysis, including the data preparation steps, number of individuals, populations and species included, analyses and the number of SNPs.

| **Dataset name** | **Data preparation steps after SNP calling** | **Total *n*** | **Populations** | **Analyses** | **No. SNPs** |
| --- | --- | --- | --- | --- | --- |
| Hard call | Removal of unplaced scaffolds | 45 | Z. lateralis (Norfolk Island)  Z. lateralis (New Zealand)  Z. lateralis x tenuirostris  Z. lateralis x albogularis  Z. tenuirostris  Z. albogularis  Z. borbonicus | Dsuite  ABBA-BABA statistics in sliding windows  Twisst2 | 1,673,542 |
| Hard call (filtered) | Removal of unplaced scaffolds  Removal of sex chromosomes  HW-pruning  LD-pruning | 45 | Z. lateralis (Norfolk Island)  Z. lateralis (New Zealand)  Z. lateralis x tenuirostris  Z. lateralis x albogularis  Z. tenuirostris  Z. albogularis  Z. borbonicus | HyDe  PCA  ADMIXTURE | 371,918 |
| Norfolk Island hard call | Removal of unplaced scaffolds  Removal of sex chromosomes  HW-pruning | 32 | Z. lateralis (Norfolk Island)  Z. tenuirostris  Z. albogularis  Z. lateralis x tenuirostris  Z. lateralis x albogularis | KING relatedness | 1,669,648 |
| Norfolk Island hard call (filtered) | Removal of unplaced scaffolds  Removal of sex chromosomes  HW-pruning  LD-pruning | 32 | Z. lateralis (Norfolk Island)  Z. tenuirostris  Z. albogularis  Z. lateralis x tenuirostris  Z. lateralis x albogularis | PCA  ADMIXTURE  TriangulaR | 224,573 |
| Norfolk Island hard call (thinned) | Removal of unplaced scaffolds  Removal of sex chromosomes  HW-pruning  LD-pruning  Thinning to 100,000 SNPs | 32 | Z. lateralis (Norfolk Island)  Z. lateralis x tenuirostris  Z. lateralis x albogularis  Z. tenuirostris  Z. albogularis  Z. lateralis x tenuirostris  Z. lateralis x albogularis | IQTREE | 100,000 |
| Norfolk Island (mitochondrial) | Filtering of SNPs to mitochondrion only | 32 | Z. lateralis (Norfolk Island)  Z. tenuirostris  Z. albogularis  Z. lateralis x tenuirostris  Z. lateralis x albogularis | Mitochondrial haplotype network with PopArt | NA (entire alignment of 29kb used) |

**Table S3**: the full results of the genomic sexing.

| **Individual** | **Source** | **W:Z Ratio** | **Inferred sex** |
| --- | --- | --- | --- |
| AEB01_L0001 | Museum | 0.098 | Male |
| AEB01_L0002 | Museum | 0.072 | Male |
| AEB01_L0003 | Museum | 0.066 | Male |
| AEB01_L0004 | Museum | 0.066 | Male |
| AEB01_L0005 | Museum | 0.073 | Male |
| AEB01_L0006 | Museum | 0.065 | Male |
| AEB01_L0007 | Museum | 0.143 | Female (degraded W) |
| AEB01_L0008 | Museum | 0.466 | Female |
| AEB01_L0009 | Museum | 0.171 | Female (degraded W) |
| AEB01_L0010 | Museum | 0.796 | Female |
| N101 | Blood sample | 0.474 | Female |
| N101 | Blood sample | 0.474 | Female |
| N102 | Blood sample | 0.059 | Male |
| N102 | Blood sample | 0.059 | Male |
| N103 | Blood sample | 0.049 | Male |
| N103 | Blood sample | 0.049 | Male |
| N104 | Blood sample | 0.049 | Male |
| N104 | Blood sample | 0.049 | Male |
| N108 | Blood sample | 0.401 | Female |
| N108 | Blood sample | 0.401 | Female |
| N110 | Blood sample | 0.478 | Female |
| N110 | Blood sample | 0.478 | Female |
| N111 | Blood sample | 0.053 | Male |
| N111 | Blood sample | 0.053 | Male |
| N114 | Blood sample | 0.055 | Male |
| N114 | Blood sample | 0.055 | Male |
| N121 | Blood sample | 0.036 | Male |
| N121 | Blood sample | 0.036 | Male |
| N122 | Blood sample | 0.048 | Male |
| N122 | Blood sample | 0.048 | Male |
| N123 | Blood sample | 0.476 | Female |
| N123 | Blood sample | 0.476 | Female |
| N126 | Blood sample | 0.433 | Female |
| N126 | Blood sample | 0.433 | Female |
| N128 | Blood sample | 0.049 | Male |
| N128 | Blood sample | 0.049 | Male |
| N129 | Blood sample | 0.043 | Male |
| N129 | Blood sample | 0.043 | Male |
| N130 | Blood sample | 0.048 | Male |
| N130 | Blood sample | 0.048 | Male |
| NOR124 | Blood sample | 0.525 | Female |
| NOR124 | Blood sample | 0.525 | Female |
| NOR125 | Blood sample | 0.528 | Female |
| NOR125 | Blood sample | 0.528 | Female |
| NOR132 | Blood sample | 0.058 | Male |
| NOR132 | Blood sample | 0.058 | Male |
| NOR134 | Blood sample | 0.065 | Male |
| NOR134 | Blood sample | 0.065 | Male |
| NOR140 | Blood sample | 0.475 | Female |
| NOR140 | Blood sample | 0.475 | Female |
| NOR141 | Blood sample | 0.062 | Male |
| NOR141 | Blood sample | 0.062 | Male |
| NOR142 | Blood sample | 0.522 | Female |
| NOR142 | Blood sample | 0.522 | Female |

**Table S4**: Full results from Dsuite. Shown are the identities of the populations tested, the D-statistic (ratio of ABBA to BABA sites), Z-score and p-value, the f4-ratio (estimated proportion of admixture), the p-value for clustering (two methods implemented), and the raw numbers of ABBA and BABA sites. NZ_zlat = New Zealand Z. lateralis, NI_Zlat = Norfolk Island Z. lateralis, Zten_hist = Z. tenuirostris (historical), Zalb = Z. albogularis. Comparisons reported in the text highlighted in **bold**.

| P1 | P2 | P3 | Dstatistic | Z-score | p-value | f4-ratio | clustering_sensitive | clustering_robust | BBAA | ABBA | BABA |
| --- | --- | --- | --- | --- | --- | --- | --- | --- | --- | --- | --- |
| **nz_zlat** | **ni_zlat** | **zalb** | **0.016475** | **12.3993** | **2.30E-16** | **0.013183** | **2.10E-14** | **0.927138** | **183659** | **78819.2** | **76264.1** |
| ni_zlat | zlat_hist | nz_zlat | 0.025955 | 6.08003 | 1.20E-09 | 0.273882 | 4.76E-13 | 5.59E-06 | 106142 | 90281.2 | 85713.3 |
| nz_zlat | ni_zlat | zten | 0.026542 | 5.14893 | 2.62E-07 | 0.016327 | 5.16E-06 | 0.002104 | 187787 | 79502.2 | 75391.1 |
| **nz_zlat** | **ni_zlat** | **zten_hist** | **0.025821** | **5.08705** | **3.64E-07** | **0.013021** | **6.84E-12** | **0.023944** | **186281** | **78817.6** | **74849.8** |
| ni_zlat | zlat_hist | zalb | 0.032153 | 5.82934 | 5.56E-09 | 0.024337 | 2.30E-16 | 2.92E-05 | 195326 | 74713.5 | 70058.7 |
| zten | zalb | ni_zlat | 0.017007 | 2.51183 | 0.012011 | 0.025623 | 1.53E-10 | 0.000954 | 136026 | 103028 | 99582.5 |
| zten_hist | zalb | ni_zlat | 0.013091 | 2.08303 | 0.037248 | 0.019637 | 0.000111 | 0.002487 | 135778 | 101553 | 98928.2 |
| zlat_hist | ni_zlat | zten | 0.026355 | 4.36678 | 1.26E-05 | 0.014965 | 2.30E-16 | 0.009299 | 201984 | 73270.1 | 69507.2 |
| zlat_hist | ni_zlat | zten_hist | 0.022303 | 3.55495 | 0.000378 | 0.010396 | 2.30E-16 | 0.135145 | 200297 | 72405.4 | 69246.1 |
| zten | zten_hist | ni_zlat | 0.011846 | 4.35205 | 1.35E-05 | 0.006108 | 2.30E-16 | 0.194931 | 269977 | 35080.9 | 34259.5 |
| nz_zlat | zlat_hist | zalb | 0.047586 | 8.63772 | 2.30E-16 | 0.037199 | 2.30E-16 | 1.87E-05 | 184114 | 79361.6 | 72151.7 |
| zten | zalb | nz_zlat | 0.025109 | 5.06053 | 4.18E-07 | 0.03876 | 1.34E-14 | 0.137868 | 137656 | 102103 | 97100.9 |
| zten_hist | zalb | nz_zlat | 0.020474 | 4.45553 | 8.37E-06 | 0.03152 | 3.23E-07 | 0.04909 | 137393 | 100612 | 96575.1 |
| nz_zlat | zlat_hist | zten | 0.002348 | 0.975599 | 0.329263 | 0.001383 | 2.30E-16 | 3.41E-06 | 190941 | 74325.3 | 73977.1 |
| nz_zlat | zlat_hist | zten_hist | 0.005493 | 1.97584 | 0.048173 | 0.002653 | 2.30E-16 | 0.000392 | 189192 | 74001 | 73192.5 |
| zten | zten_hist | nz_zlat | 0.014139 | 5.98027 | 2.23E-09 | 0.007476 | 2.30E-16 | 0.142574 | 273462 | 34597.3 | 33632.6 |
| zten | zalb | zlat_hist | 0.058612 | 9.92786 | 2.30E-16 | 0.042464 | 3.04E-06 | 0.015834 | 135477 | 107134 | 95270.9 |
| zten_hist | zalb | zlat_hist | 0.05207 | 8.93101 | 2.30E-16 | 0.037555 | 1.49E-06 | 0.029193 | 135025 | 105455 | 95016.2 |
| zten | zten_hist | zalb | 0.017041 | 5.8271 | 5.64E-09 | 0.007755 | 2.30E-16 | 0.41065 | 234674 | 36626.8 | 35399.4 |
| zten | zten_hist | zlat_hist | 0.020858 | 5.8727 | 4.29E-09 | 0.005101 | 2.30E-16 | 0.72003 | 272928 | 34872 | 33447.1 |

**Table S5:** Log-likelihood increase based on *k* parameter for ADMIXTURE.

| **k** | **Log likelihood** |
| --- | --- |
| 1 | -6480688 |
| 2 | -5718658 |
| 3 | -5365028 |
| 4 | -5190637 |
| 5 | -5054403 |
| 6 | -4889419 |
| 7 | -4732590 |
| 8 | -4581786 |
| 9 | -4506995 |
| 10 | -4356721 |

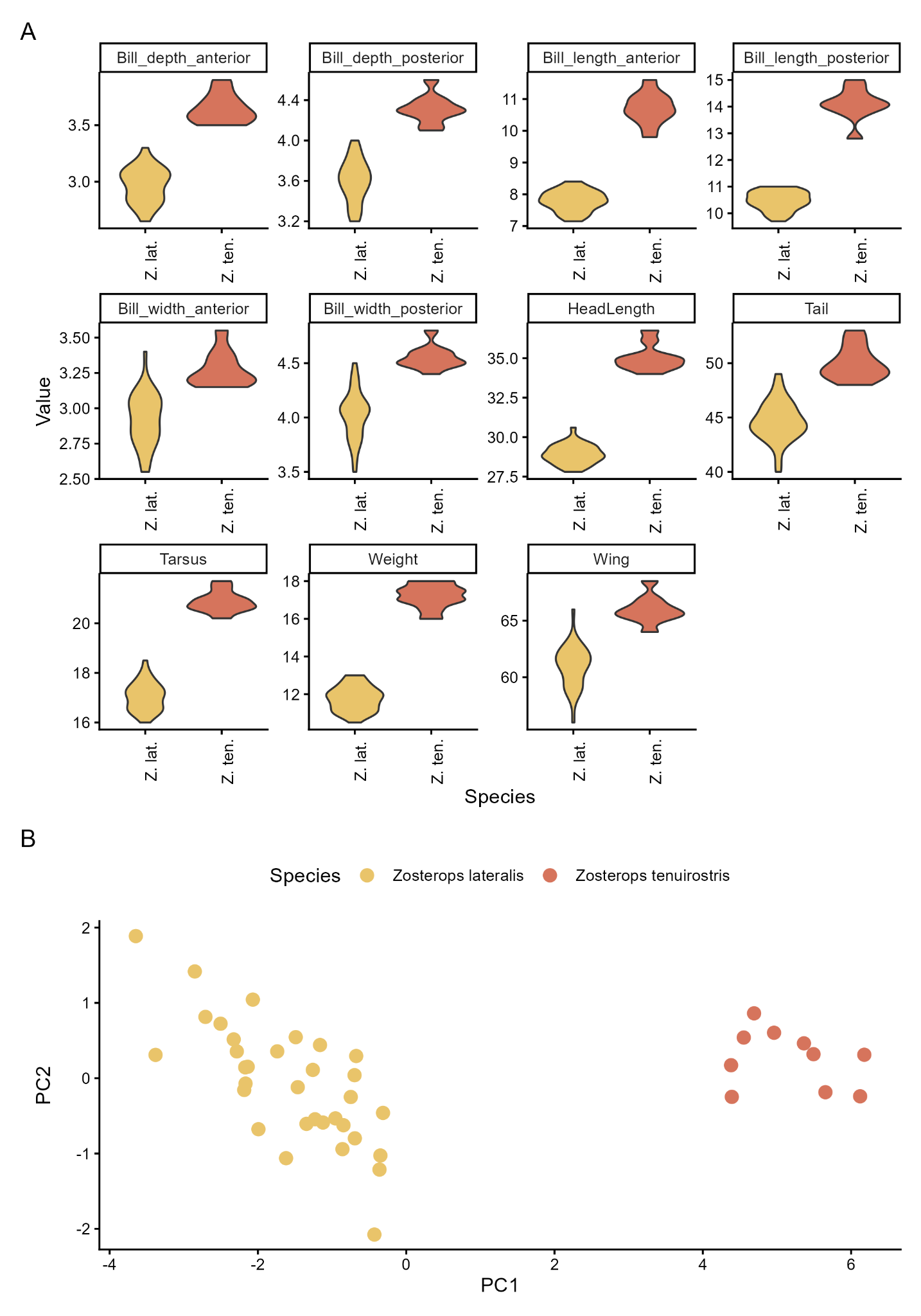

**Figure S1. A)** Morphological measurements from *Z. lateralis* and *Z. tenuirostris* taken in 1998. All values are in millimetres except for weight which is in grams. For all measurements, the distributions were significantly different between species (t-test, p-values all < 0.001). B) PCA based on results in A) showing clear separation across PC1 between species.

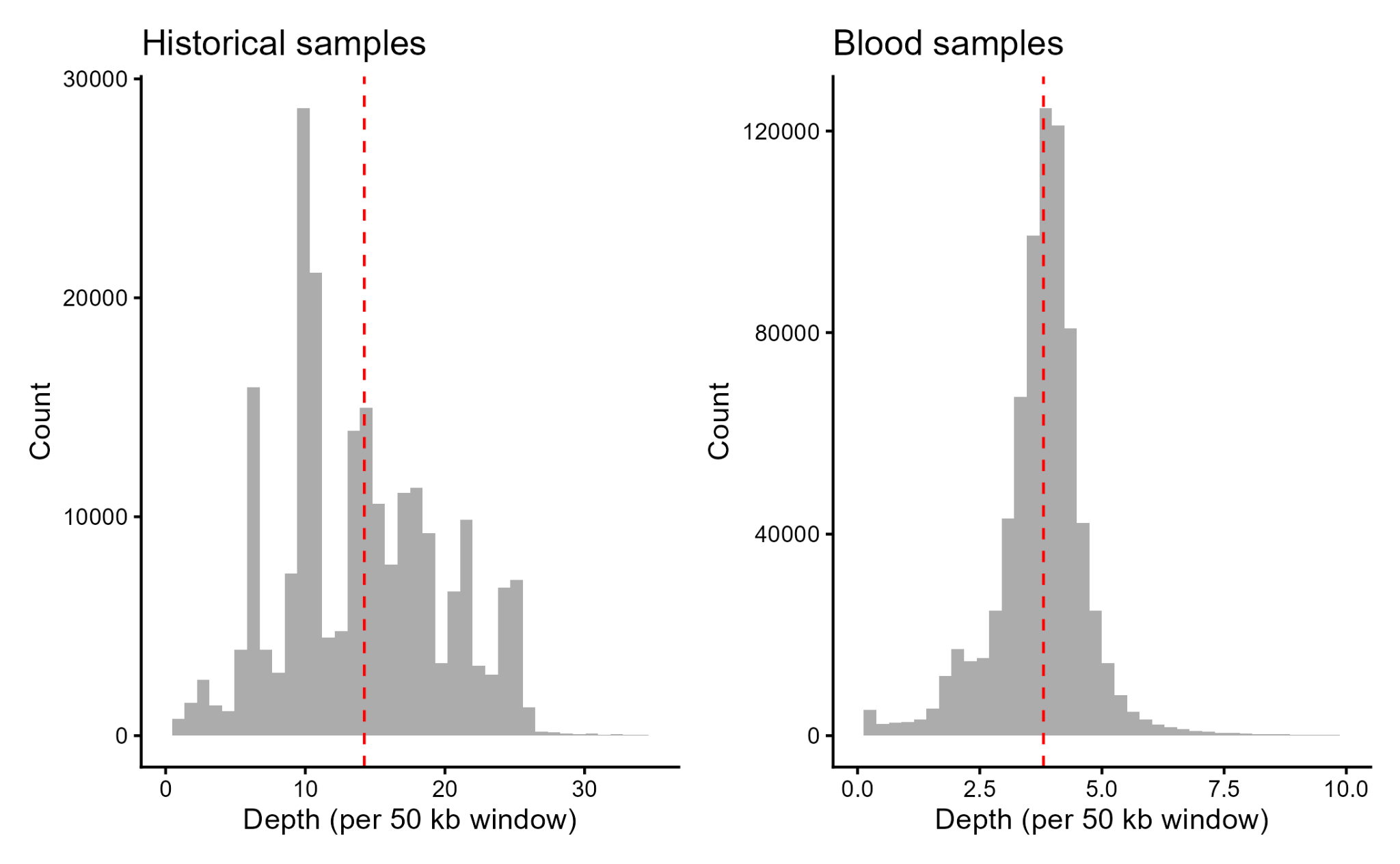

**Figure S2**. Pre-filtering depth calculated in 50kb windows and pooled across historical (high-coverage) and blood (low-coverage) samples. Red dashed vertical line indicates the mean raw coverage (14.2X for historical samples and 3.8X for blood samples).

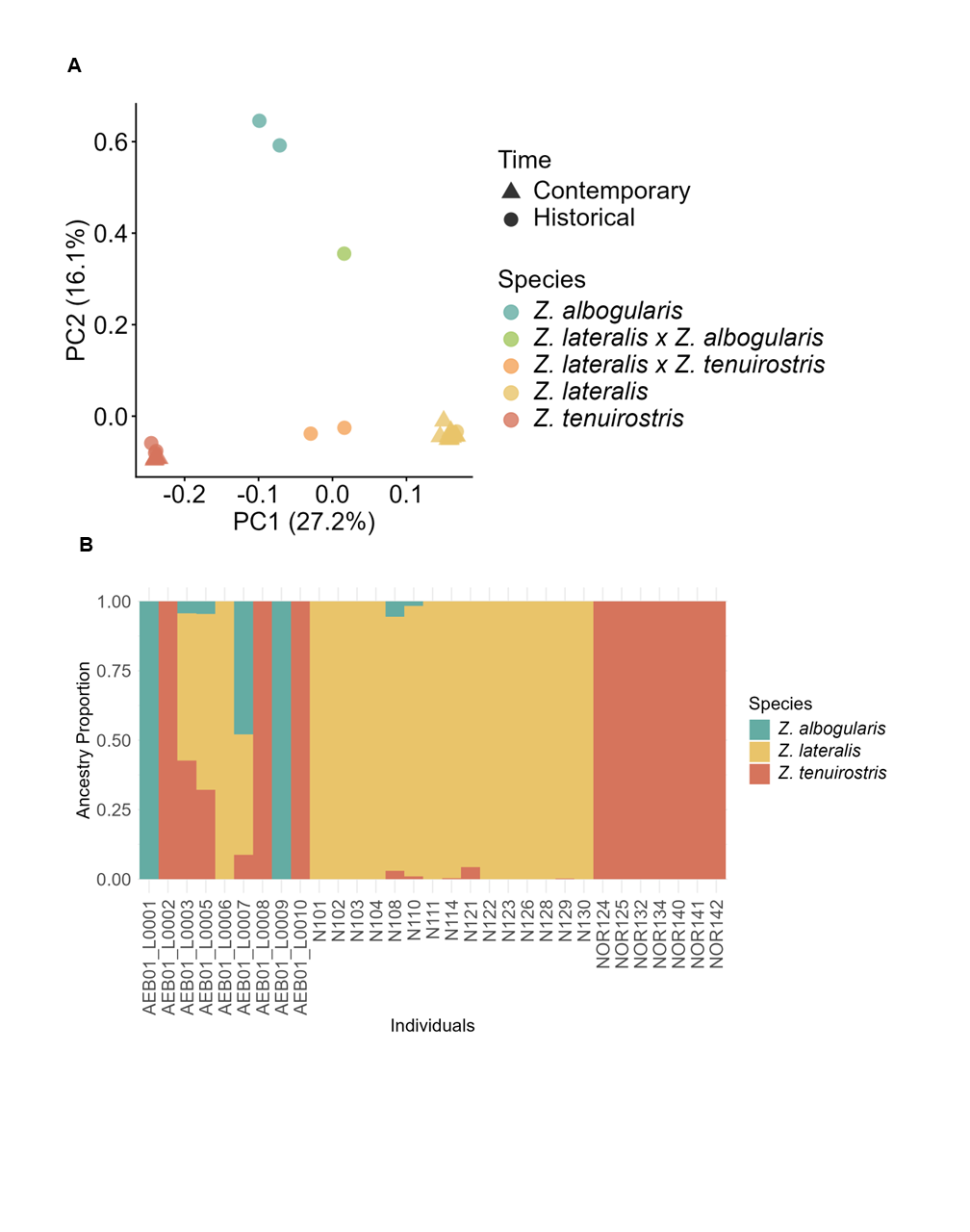

**Figure S3**. PCA (A) and ADMIXTURE (B) results (performed on hard call dataset) with related individuals removed. Here we removed AEB01_L0004, a *Z. lateralis-Z. tenuirostris* hybrid, which was likely a third-degree relative of AEB01_L0003.

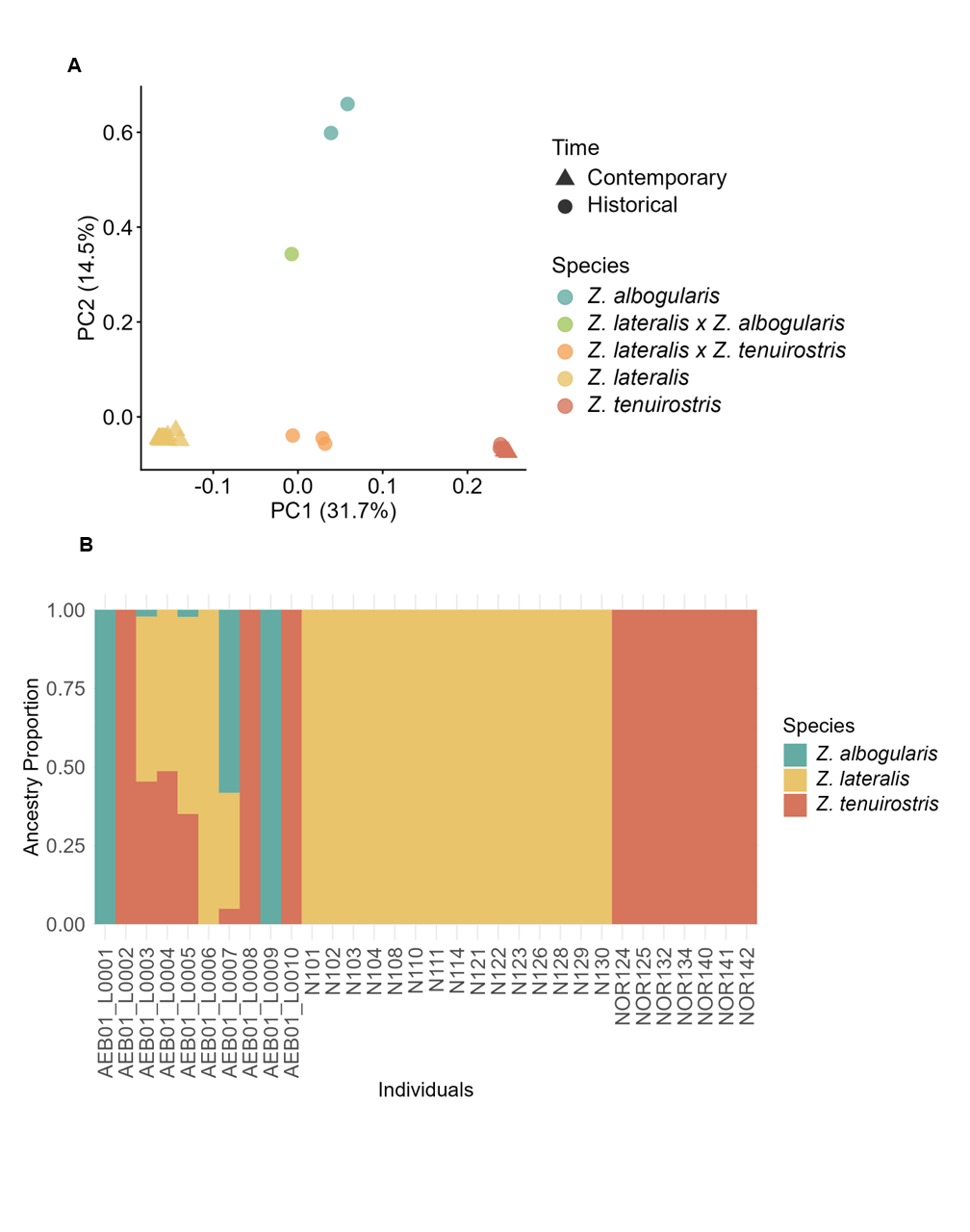

**Figure S4.** Population structure results produced within a genotype likelihood framework (ANGSD). A) PCAngsd B) NGSadmix.

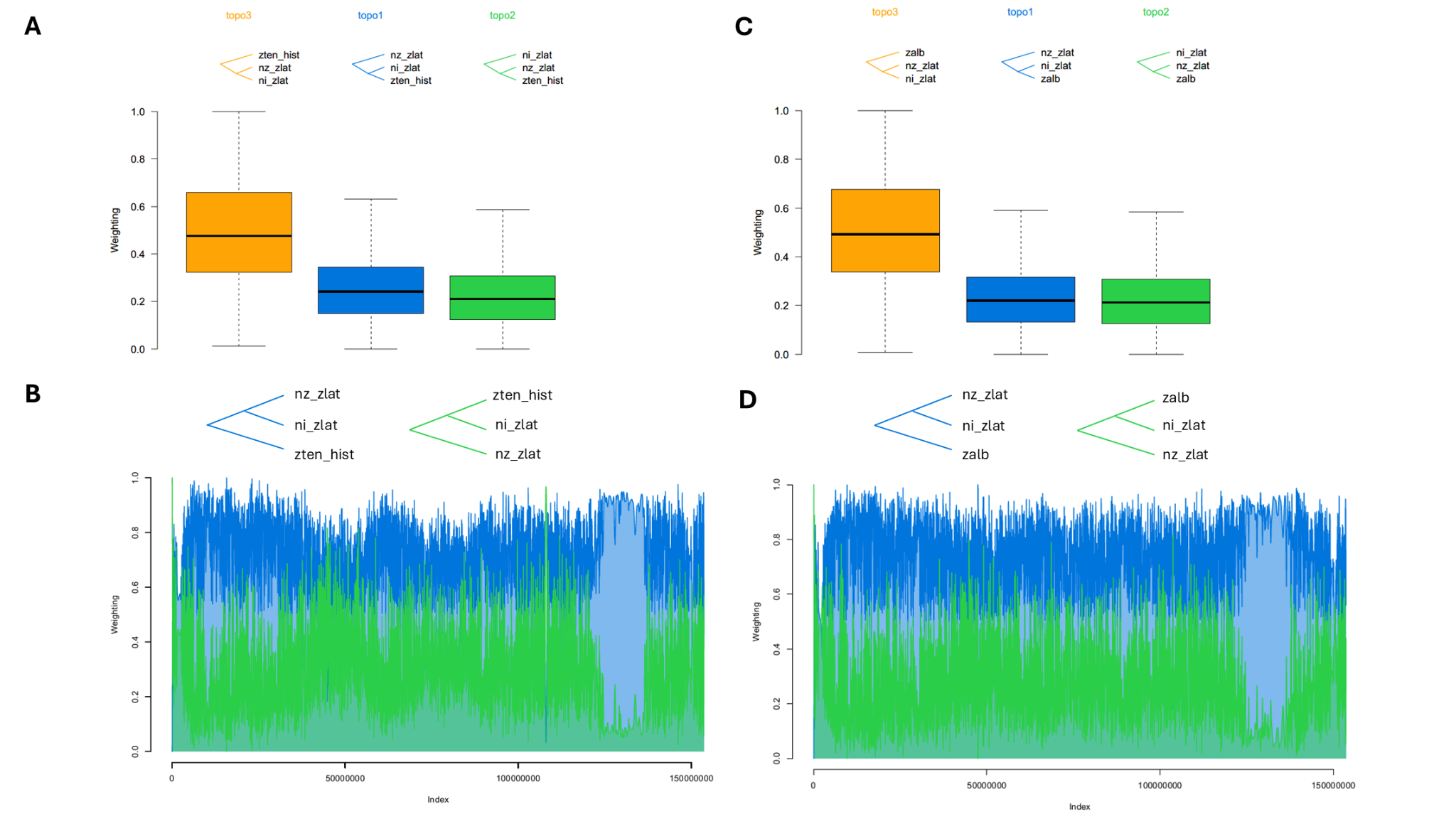

**Figure S5**. Results from Twisst2 topology weighting software. Panels A and B refer to the comparison involving *Z. tenuirostris* (zten_hist) and panels C and D refer to the comparison involving *Z. albogularis* (zalb). Panels A and C show the overall genome-wide topology weighting with topologies shown. As expected, in both comparisons, topology 3 (orange) shows the highest weighting due to the closer phylogenetic relationship of the two *Z. lateralis* populations (nz_zlat = New Zealand *Z. lateralis*; ni_zlat = Norfolk Island *Z. lateralis*). Topology 1 (blue) is the next best-supported, followed by Topology 2 (green). Panels B and D show the topology weighting genome scan with colours corresponding to the topologies shown. Only chromosome 1 is shown for clarity.
